# Rethinking the role of lipids in lager yeast cells during beer fermentation from a transcriptome and systems biology perspective

**DOI:** 10.1101/2020.01.28.922898

**Authors:** Diego Bonatto

## Abstract

Brewing lager yeast (*Saccharomyces pastorianus*) is exposed to stressful conditions during beer fermentation, including ethanol toxicity. In response to ethanol toxicity, various biological mechanisms are modulated, including lipid biosynthesis. It is well known that during beer fermentation, the composition of yeast membranes changes in response to ethanol toxicity, making it less fluid and permeable. Additionally, neutral lipids and lipid droplets (LDs) are produced in response to ethanol toxicity. LDs are membranous organelles that transport lipids and proteins, acting as hubs for inter-organellar communication and modulating the activity of mechanisms necessary for ethanol tolerance, such as proteostasis and autophagy. Unfortunately, little is known about the interplay between autophagy, lipid metabolism, and proteostasis (ALP) in lager cells during beer fermentation. Therefore, transcriptome analyses using publicly available DNA microarray data obtained from lager yeast cells were used to identify all the ALP-associated genes that were upregulated during beer fermentation compared to yeast biomass propagation. Thereafter, a top-down systems biology analysis was applied, involving constructing an ALP-associated shortest-pathway protein–protein interaction network (ALP network), identifying important nodes and communities within the ALP network, and identifying the overrepresented biological processes and cellular components using a Gene Ontology (GO) analysis. The transcriptome analyses indicated the upregulation of 204 non-redundant ALP-associated genes during beer fermentation, whose respective proteins interact in the shortest-pathway ALP network. Thirteen communities were selected from the ALP network, and they were associated with multiple overrepresented GO biological processes and cellular components, such as mitophagy, cytoplasm-to-vacuole transport, piecemeal microautophagy of the nucleus, endoplasmic reticulum (ER) stress, ergosterol and lipid biosynthesis, LDs, ER membrane, and phagophore assembly. These results indicate that ethanol tolerance in lager yeasts could be due to the modulation of proteostasis and various forms of autophagy by lipid biosynthesis and LDs, thus highlighting the importance of lipids for beer fermentation.

## 1. Introduction

Nowadays, high gravity (HG) and very high gravity (VHG) fermentation technologies are often used in the brewing industry for beer production. This reduces the consumption of water during the brewing process and increases the ethanol yield, thus maximizing the brewing efficiency and reducing the production costs and energy demand (Puligundla et al., 2019, 2011). However, the accumulation of large amounts of ethanol (>5% v/v) due to the fermentation of HG/VHG wort drastically alters the yeast’s physiology (Hallsworth, 1998) and promotes an ethanol stress response that induces molecular mechanisms associated with the heat shock response (Odumeru et al., 1992; Piper, 1995).

The ability of different yeast strains to cope with the toxic effects of ethanol depends on the modulation of cell membrane fluidity by alteration of the ratio of incorporated saturated and unsaturated fatty acids and the ergosterol content (Ding et al., 2009). It is well established that membrane-associated lipids have a strong influence in beer brewing, affecting the fermentative capacity and ethanol tolerance of *Saccharomyces cerevisiae* (ale yeast) and *Saccharomyces pastorianus* (lager yeast) (Ahvenainen, 1982; Mishra and Kaur, 1991). In wine yeast strains, the high concentration of ergosterol in the cell membrane promotes ethanol tolerance by decreasing membrane fluidity (Aguilera et al., 2006); however, the increased levels of unsaturated fatty acids in the cell membrane increase the membrane fluidity (Alexandre et al., 1994). It was found that yeast strains that are more ethanol tolerant incorporate long-chain fatty acids (C_18:0_ and C_18:1_) compared with strains that are less ethanol tolerant (Chi and Arneborg, 1999). Additionally, high concentrations of ethanol induce the fluidification and thinning of membranes, along with changing the activity and aggregation of membrane-associated proteins (Thibault et al., 2012).

Unfortunately, little is known about how lipids modulate various biological mechanisms in yeast cells during beer fermentation besides affecting membrane structure and/or permeability. However, it is known that membrane fluidification by ethanol can activate the endoplasmic reticulum (ER)- linked unfolded protein response (UPR) (Navarro-Tapia et al., 2018) and lipids may have other roles in proteostasis, such as the removal of unfolded proteins from the ER by lipid droplets (LDs) (Vevea et al., 2015). LDs are important and highly dynamic cytoplasmic organelles that connect different parts of the cell, including the ER (Jacquier et al., 2011), mitochondria (Pu et al., 2011), peroxisomes (Kohlwein et al., 2013), and vacuoles (Barbosa et al., 2015). Interestingly, ER stress induces the formation of LDs (Fei et al., 2009) and stimulates lipid biosynthesis, which are associated with ER membrane expansion during UPR (Cox et al., 1997; Schuck et al., 2009). Additionally, lipid biosynthesis coordinates the proteotoxic response of both mitochondria and the cytosol (Kim et al., 2016). Thus, it can be hypothesized that LD-associated processes and lipid biosynthesis are integrative processes necessary for proteostasis in different organelles. In fact, regulation of inter-organellar proteostasis is an important mechanism underlying stress tolerance and it was recently shown that beer fermentation in lager yeast cells promotes a so-called “inter-organellar/cross-organellar communication/response” (CORE mechanism). This involves a series of signaling-associated protein networks that regulate inter-organellar proteostasis, which includes the ER and mitochondria UPRs, chaperone and co-chaperone activity, and *N-*glycosylation quality control pathway proteins (Telini et al., 2020). A major aspect of the inter-organellar proteostasis mechanism induced by ethanol stress is the coordination and/or activation of organellar-linked microautophagy responses, such as mitophagy (Carmona-Gutierrez et al., 2012) and lipophagy of LDs (Vevea et al., 2015). Moreover, macroautophagy can also be induced by reactive oxygen species (ROS) generated by damaged mitochondria due to ethanol stress (Jing et al., 2018).

Thus, the purpose of this study was to evaluate how lipid metabolism interacts with different mechanisms linked to inter-organellar proteostasis and autophagy in lager beer yeast during beer fermentation. Publicly available DNA microarray gene expression datasets obtained from lager beer yeast at various time points during industrial beer fermentation and yeast biomass propagation were selected and two transcriptome analyses were performed. The differentially expressed genes that were upregulated in both analyses (Pan-DEGs) were used to generate a protein–protein interaction (PPI) network. This was followed by local and global topological analyses as well as a gene ontology (GO) analysis of major clusters within the PPI network. The results of the transcriptome and PPI network analyses indicated cross-communication between various pathways linked to inter-organelle autophagy, lipid metabolism, and proteostasis (ALP) during lager beer fermentation.

## 2. Material and methods

### 2.1 DNA microarray gene expression dataset selection and analysis

DNA microarray gene expression datasets (GSE9423, GSE10205, and GSE16376) containing transcriptome data obtained from the lager yeast strain CB11 (*Saccharomyces pastorianus*) at various time points during industrial beer fermentation and yeast biomass propagation were selected from the Gene Expression Omnibus (GEO) database (http://www.ncbi.nlm.nih.gov/gds) (Table S1). The GSE9423 dataset contains the transcriptome data of strain CB11 during both beer fermentation (time points: 8, 30, and 60 h) and yeast biomass propagation (time points: 0, 8, and 30 h) (Gibson et al., 2008), and the analysis based on these data was designated the “single-analysis”. The GSE10205 and GSE16376 datasets contain the transcriptome data of strain CB11 during beer fermentation (time points: 8, 30, 60, 80, and 102 h) and yeast biomass propagation (time points: 0, 4, 8, and 30 h), respectively. They were combined for an analysis designated the “meta-analysis” (Suppl. Fig. 1). It is important to note that both the GSE10205 and GSE16376 datasets evaluated the same lager yeast strain (CB11) in identical fermentation and propagation conditions used in GSE9423 dataset. Moreover, the number of high-throughput (RNA-seq) studies involving *S. pastorianus* in conditions of yeast biomass propagation and beer fermentation are virtually nonexistent until now. In this sense, the presence of a high number of orthologous sequences and the closest phylogeny of *Saccharomyces cerevisiae* and *Saccharomyces eubayanus*, which composes the hybrid genome of *S. pastorianus*, allows to apply DNA microarray platform available for *S. cerevisiae* for *S. pastorianus* transcriptome analysis. Horinouchi et al. (2010) discuss the use of a tailor-made DNA microarray for *S. pastorianus* and show a strong correlation between the expression levels of *S. cerevisiae* and *S. bayanus* orthologous genes during fermentation (Horinouchi et al., 2010).

The transcriptome analyses were performed using the R platform (https://www.r-project.org) with various packages (Suppl. Fig. 1). For data matrix importing, processing, and array quality analysis, the GEOquery, affy, and arrayQualityMetrics packages, respectively, were employed (Davis and Meltzer, 2007; Gautier et al., 2004; Kauffmann et al., 2009). DEG analysis was performed using the limma package (Ritchie et al., 2015). The false discovery rate (FDR) algorithm, implemented in the limma package (Ritchie et al., 2015), was used to assess the significance level of the DEGs. DEGs from the single-analysis (GSE9423) and meta-analysis (GSE10205 versus GSE16376) with mean |log(fold change [FC])| ≥2.0 and FDR <0.05 were selected and the ALP-associated genes were identified using *Saccharomyces cerevisiae* annotation data from the *Saccharomyces* Genome Database (https://www.yeastgenome.org). The ALP files containing the curated data used in this study are available at https://github.com/bonattod/Lipid_stress_data_analysis.git under the names “Autophagy_Table.txt”, “Lipid_metabolism_Table.txt”, and “Proteostasis_Table.txt”. For the further analyses, only upregulated DEGs (i.e., DEGs that were upregulated during beer fermentation compared to yeast biomass propagation) that were common to both the single-analysis and meta-analysis were selected. For each Pan-DEG, meta-log_2_FC ± standard deviation (SD) were calculated, and the ALP Pan-DEGs were used for ALP network design and analyses (Suppl. Fig. 1). Pan-DEGs associated with LDs in yeast were also selected using data reported by Grillitsch et al. (2011).

### 2.2 Network design and topology analyses

Initially, PPI and chemical-protein interaction (CPI) networks for *Saccharomyces cerevisiae* were designed using *S. cerevisiae* interactome data downloaded from STRING 11.0 (https://string-db.org) and STITCH 5.0 (http://stitch.embl.de), and processed in the R environment (Suppl. Fig. 1).

The *S. cerevisiae* interactome data were filtered by selecting the subscore information variables labeled “experiments” and “curated databases”, followed by the generation of a combined score from the two channels using the equation described by (von Mering et al., 2005). The resulting “String_data_2019_11.txt” and “Stitch_data_2020_5.txt” files were used for network design and topology analysis. It contains the source nodes (“Feature”), target nodes (“Feature_2”), and edge information (“combined_score”) and can be downloaded from the GitHub repository (https://github.com/bonattod/Lipid_stress_data_analysis.git). From PPI interactome network, an ALP network was obtained by selecting the shortest pathways among the Pan-DEGs using the R package igraph (Csardi and Nepusz, 2006). By its turn, the CPI network was applied to select the shortest pathways among LDs-associated proteins and lipid molecules in yeast (Suppl. Fig. 1) and to generate a multilayered network containing both transcriptome and proteome data (LDP network). In this sense, the proteome data was obtained from (Casanovas et al., 2015) and used to evaluate the expression of LDs-associated proteins during the fermentation-to-respiration transition (Suppl. Fig. 1). Chemical data of interacting lipid molecules was obtained from Lipid Maps^®^ Lipidomics Gateway (https://lipidmaps.org/). ALP and LDP network were visualized in Cytoscape 3.7.2 (Shannon et al., 2003) using the RCyc3 package (Gustavsen et al., 2019) (Suppl. Fig. 1).

Regarding the centrality analysis of ALP network (Suppl. Fig. 1), the node degree and betweenness values were calculated using the R package igraph (Csardi and Nepusz, 2006). Node degree indicates the number of connections that a specific node has, while betweenness indicates the number of shortest paths that pass through a specific node. All nodes that had a degree value above the mean for the network were designated “hubs”, while all nodes that had a betweenness value above the mean of the network were designated “bottlenecks” (Yu et al., 2007). Finally, the node degree and betweenness values were used to group the nodes into four major groups: (i) hub-bottleneck (HB), (ii) nonhub-bottleneck (NHB), (iii) hub-nonbottleneck (HNB), and (iv) nonhub-nonbottleneck (NHNB). The HB group represents all nodes that potentially control the flow of information through the network and act as key regulators in the cell (Yu et al., 2007).

Community analysis of ALP network was performed in the R environment using the walktrap community (WTC) finding algorithm described by (Pons and Latapy, 2005), and the analysis was fully implemented using the igraph package (Csardi and Nepusz, 2006). Communities were selected on the basis of two criteria: (i) presence of HB nodes and (ii) presence of Pan-DEGs (Suppl. Fig. 1). The selected clusters were visualized in Cytoscape 3.7.2 (Shannon et al., 2003) using the R package RCyc3 (Gustavsen et al., 2019) (Suppl. Fig. 1).

### 2.3 GO analysis

The GO biological process and cellular component categories associated with the selected communities from the ALP network were determined using the R package clusterProfile (Yu et al., 2012) (Suppl. Fig. 1). The degree of functional enrichment for each biological process and cellular component category was quantitatively assessed (*p* < 0.01) using a hypergeometric distribution. Multiple testing correction was also implemented using the FDR algorithm (Benjamini and Hochberg, 1995) with a significance level of *p* < 0.05. Semantic comparison of the biological processes and cellular components associated with the node clusters was conducted using the R package GOSemSim (Yu et al., 2010) (Suppl. Fig. 1) using FDR < 0.01 and *q* < 0.05. Heatmaps combining GO categories (columns) and selected clusters from the ALP network (rows) were designed using the R package ComplexHeatmap (Gu et al., 2016) (Suppl. Fig. 1), with columns and rows both being grouped using the *k-*means distance-based method (Suppl. Fig. 3 and 4).

### 2.4 Data sharing repository

All files and figures generated in this study can be freely downloaded from https://github.com/bonattod/Lipid_stress_data_analysis.git.

## 3. Results

### 3.1 Transcriptome single- and meta-analyses of ALP-associated genes in lager yeast strain CB11 during beer fermentation

The initial comparison of upregulated and downregulated DEGs in strain CB11 (during industrial beer fermentation compared to yeast biomass propagation) identified in the GSE9423 dataset analysis (single-analysis; Suppl. Fig. 1) and the GSE10205 versus GSE16376 dataset analysis (meta-analysis; Suppl. Fig. 1) indicated similar patterns of upregulated and downregulated DEGs (Suppl. Fig. 2A and B). There were a total of 5,134 upregulated and 4,954 downregulated DEGs in the single-analysis (Suppl. Fig. 2A and Table S2), while there were a total of 10,258 upregulated and 9,342 downregulated DEGs in the meta-analysis (Suppl. Fig. 2A and Table S2). It should be noted that the high frequency of total upregulated and downregulated DEGs was due to gene redundancy across the various comparisons in both the single- and meta-analyses. After removing the redundant genes from both analyses, there were 1,315 upregulated and 1,209 downregulated non-redundant (unique) DEGs in the single-analysis, and 1,727 upregulated and 1,502 downregulated non-redundant DEGs in the meta-analysis (Suppl. Fig. 2B).

After the transcriptome single- and meta-analyses were conducted (Table S2), the next step was to evaluate the expression profile of ALP-associated genes. The gene information in the *Saccharomyces* Genome Database regarding ALP mechanisms (Suppl. Fig. 1) was used to select the ALP-associated DEGs. The total and non-redundant frequencies of upregulated ALP DEGs were higher than the frequencies of downregulated ALP DEGs in both the single-analysis and meta-analysis (Suppl. Fig. 2C and D). Regarding the upregulated ALP DEGs (for both redundant and non-redundant DEGs), a high frequency of lipid metabolism-associated genes was observed in both transcriptome analyses, followed by proteostasis-associated genes and then autophagy-associated genes (Suppl. Fig. 2C and D). Interestingly, the frequencies of redundant and non-redundant downregulated ALP DEGs were similar across the transcriptome analyses for all three ALP processes (Suppl. Fig. 2C and D).

This initial transcriptome data evaluation was followed by a specific analysis of the absolute a frequencies of ALP DEGs in each pairwise comparison of beer fermentation and yeast biomass propagation at various time points (Suppl. Fig. 3 and Tables S3 and S4). The absolute frequencies were low when beer fermentation was compared to early propagation time points (0 h in the single-analysis; 0 and 4 h in the meta-analysis) (Suppl. Fig. 3 and Tables S3 and S4). On the other hand, the absolute frequencies increased when beer fermentation was compared to advanced propagation time points (8 and 30 h in both the single- and meta-analyses) (Suppl. Fig. 3 and Tables S3 and S4). For the subsequent transcriptome and systems biology analyses, only the upregulated non-redundant (unique) ALP DEGs observed in all pairwise comparisons were considered.

### 3.2 ALP Pan-DEGs in lager yeast strain CB11 during beer fermentation

The frequency of non-redundant upregulated ALP DEGs for each of the three ALP processes was similar between the single-analysis and meta-analysis (Fig. 1A). Regarding autophagy, there were 27 upregulated DEGs for the single-analysis and 42 for the meta-analysis (Fig. 1A). Regarding proteostasis, there were 76 upregulated DEGs for the single-analysis and 88 for the meta-analysis (Fig. 1A). Regarding lipid metabolism, there were 121 upregulated DEGs for the single-analysis and 138 for the meta-analysis (Fig. 1A).

**Fig. 1.**
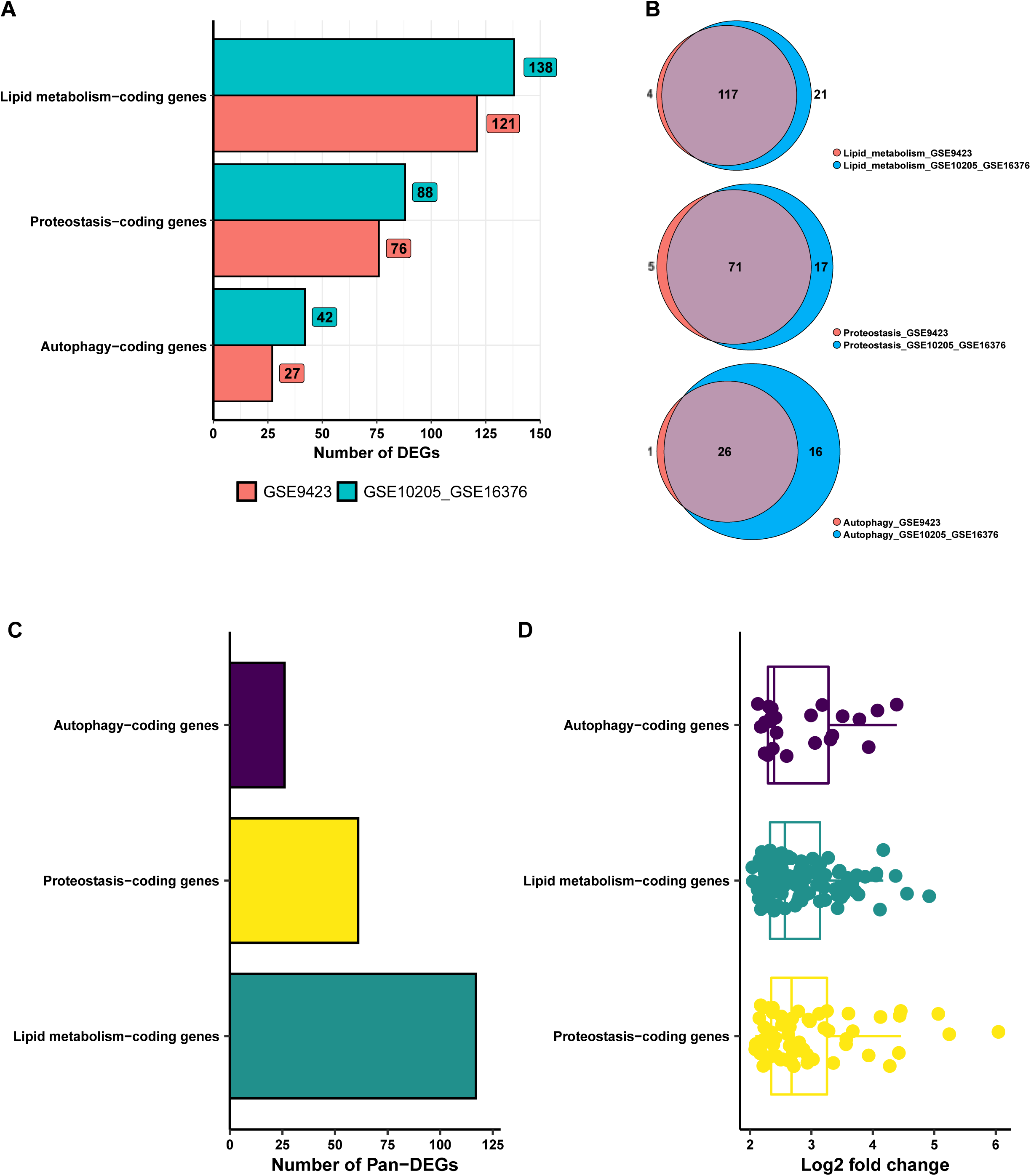
(A) Frequency of non-redundant (unique) upregulated differentially expressed genes associated with autophagy, lipid metabolism, and proteostasis (ALP DEGs) in the transcriptome single-analysis (GSE9423) and meta-analysis (GSE10205 versus GSE16376). The numbers inside the squares show the frequency of upregulated DEGs associated with a specific ALP process. The frequencies of ALP DEGs common to both transcriptome analyses (designated ALP Pan-DEGs) are indicated in the (B) Venn diagram and (C) bar chart. (D) Expression values of ALP Pan-DEGs in log_2_(fold change).

Next, the frequency of non-redundant upregulated ALP DEGs that were common to both transcriptome analyses was evaluated (Fig. 1B). There were 117, 71, and 26 common DEGs for lipid metabolism, proteostasis, and autophagy, respectively (Fig. 1B and C). These common ALP DEGs were designated ALP Pan-DEGs (Table S6) and their expression patterns were evaluated.

Regarding the expression patterns of the ALP Pan-DEGs (Fig. 1D and Table S6), the median meta-log_2_FC value was similar for autophagy (2.39), lipid metabolism (2.57), and proteostasis (2.68). Additionally, the minimum and maximum meta-log_2_FC values were similar, from 2.13 to 4.39 for autophagy, 2.09 to 4.45 for proteostasis, and 2.04 to 4.17 for lipid metabolism (Fig. 1D).

The similar expression patterns of the ALP Pan-DEGs prompted us to evaluate how the ALP Pan-DEGs are connected to each other in terms of PPIs, using a top-down systems biology approach involving *S. cerevisiae* interactome data (Suppl. Fig. 1). Moreover, the importance of the ALP Pan-DEGs in the local and global PPI network topologies was also assessed, and the overrepresented biological processes and cellular components associated with the PPI network were identified.

### 3.3 Top-down systems biology analysis of ALP Pan-DEGs

The ALP Pan-DEGs obtained from the transcriptome analyses were selected as seeds to generate a shortest-pathway PPI network using publicly available *S. cerevisiae* interactome data (Suppl. Fig. 1). The *S. cerevisiae* interactome data were filtered using only the subscore information variables named “experiments” and “curated databases” to provide evidence of interactions among proteins (as described previously in this study). This led to a PPI network composed of 5,063 nodes and 185,404 edges (see the Supplementary Files “graph_string_db_exp.txt” and “Cytoscape_data.cys” available at https://github.com/bonattod/Lipid_stress_data_analysis for more details). A subnetwork containing all the shortest pathways among the proteins encoded by the ALP Pan-DEGs was generated from the *S. cerevisiae* interactome network (Suppl. Fig. 1). This subnetwork, designated the “ALP network”, contained 1,705 nodes and 22,806 edges (Fig. 2A) and included almost all ALP Pan-DEGs with the exception of *FAT3, IZH2, IZH4, MZM1, OPI10, PPX1*, and *TMA17*, which could not be mapped using the currently available *S. cerevisiae* interactome data.

**Fig. 2.**
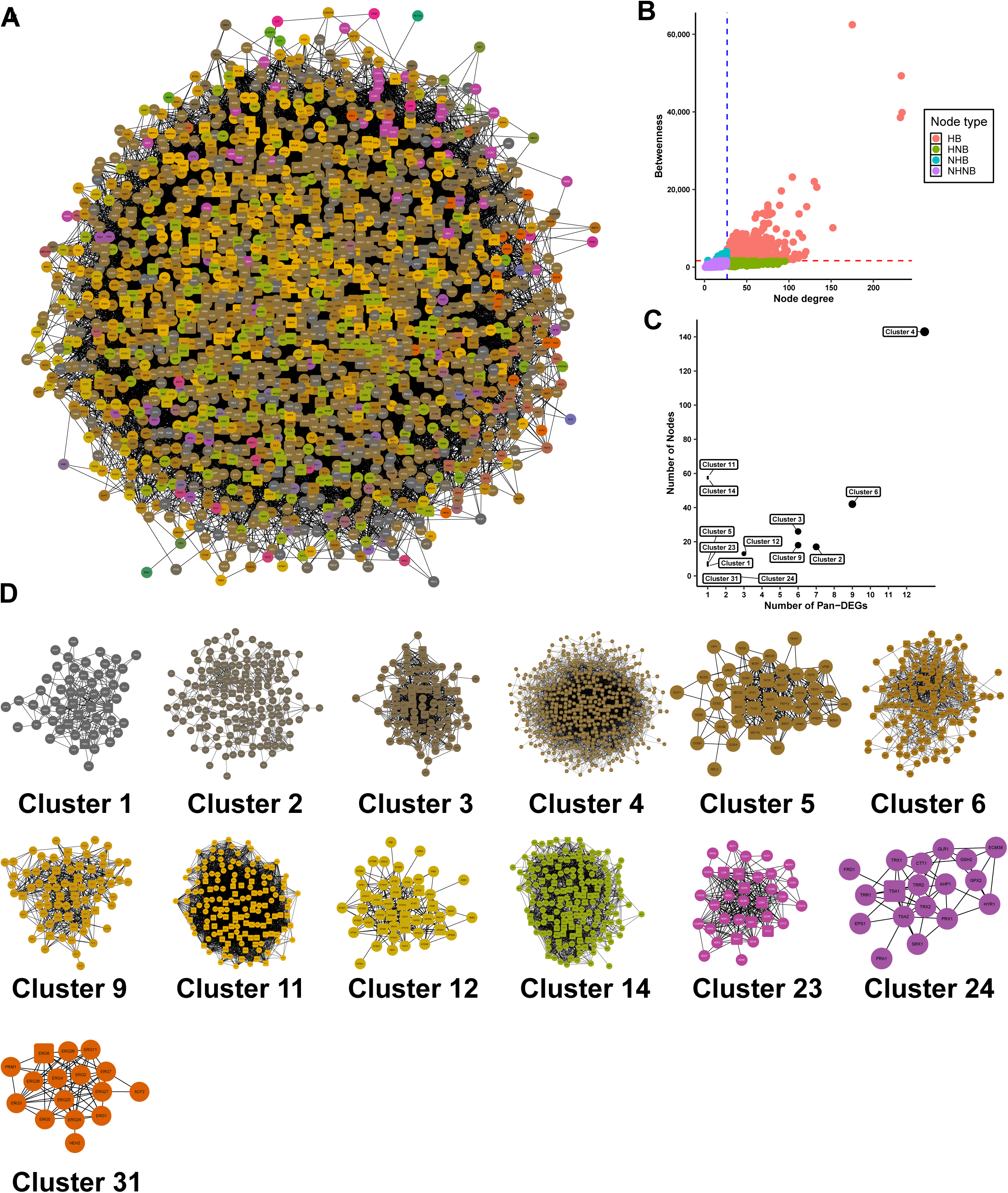
(A) Shortest-pathway protein–protein interaction (PPI) network (ALP network) based on common differentially expressed genes associated with autophagy, lipid metabolism, and proteostasis (ALP Pan-DEGs). (B) A centrality analysis was applied to this ALP network and the nodes were classified as hub-bottleneck (HB), hub-nonbottleneck (HNB), nonhub-bottleneck (NHB), or nonhub-nonbottleneck (NHNB) based on the node degree and betweenness values relative to the mean values of the network (indicated by the dashed lines). Additionally, communities/clusters of proteins in the ALP network were identified and selected based on the presence of (C) ALP Pan-DEGs (dot diameters are proportional to the number of ALP Pan-DEGs in each community/cluster) and (D) HB nodes (HB nodes are represented by square elements).

Following the generation of the ALP network, a node centrality analysis was performed to identify all nodes that exert a local influence in the network and, consequently, may have roles in ethanol stress tolerance in lager yeast cells during beer fermentation. For this purpose, two centrality parameters commonly used for PPI network analyses were selected: node degree and betweenness. The node degree evaluates the potential of a protein to connect with other proteins, thus forming functional complexes (Yu et al., 2007). All proteins with node degree values above the mean value of the network were defined as hubs (Yu et al., 2007). The betweenness evaluates the ability of a node to connect to nodes in different clusters/communities, thus serving as a “bottleneck” for biological information to traverse from one community to another (Yu et al., 2007). By combining the node degree and betweenness results, nodes that display high values for both parameters (HB nodes) can be selected (Fig. 2B). HB nodes are critical elements within a network as they concentrate the highest numbers of shortest pathways and connections with other nodes, so they are important components for signal transduction among protein clusters/communities (Yu et al., 2007). The centrality analysis of the ALP network indicated the presence of 423 HB nodes, 221 HNB nodes, 59 NHB nodes, and 1,002 NHNB nodes (Fig. 2B and Supplementary File “topologies_lipid_stress_data.txt” available at https://github.com/bonattod/Lipid_stress_data_analysis). Once the centralities in the ALP network were defined, whether they were organized into communities was determined. In general, a community can be defined as a specific network topology that contains highly connected nodes that have low degree values with respect to nodes outside the community. Moreover, communities can potentially be associated with specific biological processes (Pons and Latapy, 2005; Ravasz et al., 2002). To identify the communities in the ALP network, the WTC algorithm was applied (Pons and Latapy, 2005), which allows communities to be efficiently identified by using the random walk technique. By using the WTC algorithm, 36 communities were identified in the ALP network (see Supplementary File “topologies_lipid_stress_data.txt” available at https://github.com/bonattod/Lipid_stress_data_analysis). For further analysis, it was necessary to select the major communities in the ALP network by determining the presence of HB nodes and ALP Pan-DEGs within the communities. Thus, 13 communities were selected (Fig. 2C and D and Table S7). Cluster 4 was the largest community, with 521 nodes and 4,424 edges, and Cluster 31 was the smallest community, with 16 nodes and 69 edges (see Supplementary File “Cytoscape_data.cys” available at https://github.com/bonattod/Lipid_stress_data_analysis). Additionally, Cluster 4 contained the largest number of ALP Pan-DEGs, while Cluster 24 contained only one ALP Pan-DEG (Suppl. Fig. 4 and Table S7).

Following the community identification, a GO analysis was applied to each of the 13 selected clusters in order to identify the major overrepresented biological process categories (Fig. 3 and Table S8) and cellular component categories (Fig. 4 and Table S9). Regarding biological processes, three cluster groups and seven biological process groups were identified by applying a *k-*means distance-based method (Fig. 3). Cluster group 1 (clusters 3, 23, and 24) was associated with major biological processes concerning mitochondria structure and organization (Fig. 3), while cluster group 2 (clusters 2, 9, and 31) was associated with lipid, ergosterol, and alcohol metabolism (Fig. 3). Both cluster groups 1 and 2 also contained proteins involved in the oxidation-reduction process (Fig. 3). On the other hand, cluster group 3 (clusters 1, 4, 5, 6, 12, 10, and 14) was associated with autophagy and autophagosome assembly, response to ER stress and protein folding, piecemeal microautophagy of the nucleus (PMN) and mitochondria autophagy, vesicle-mediated transport, and other biological processes (Fig. 3). Next, the major cellular components associated with the clusters were identified. Again, three cluster groups were observed (Fig. 4). Cluster group 1 (clusters 1, 4, and 31) mainly contained proteins associated with ER membrane and LDs, while cluster group 2 (clusters 2, 3, 5, 9, and 23) contained proteins found in the mitochondria envelope and matrix, Golgi apparatus, membrane protein complexes, and organelles such as peroxisomes (Fig. 4). Finally, cluster group 3 (clusters 8 and 12) contained proteins that are mainly found in cytoplasmic vesicles and at phagophore assembly sites (Fig. 4).

**Fig. 3.**
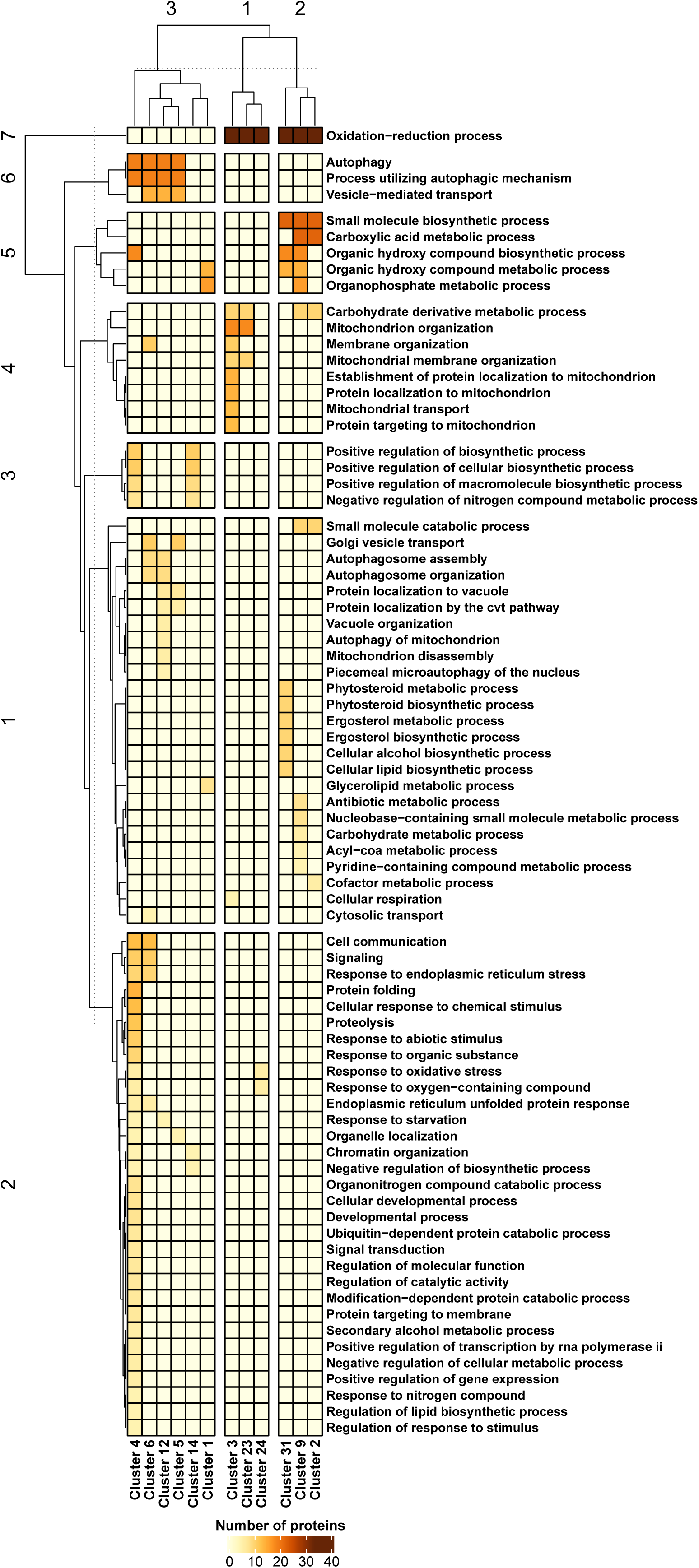
Heatmap showing the clustered biological processes based on the gene ontology (GO) analysis of autophagy, lipid metabolism, and proteostasis (ALP) network-associated communities/clusters. Heatmap rows and columns were grouped using the *k-*means distance-based method. Horizontal and vertical dotted lines indicate the cut-off point used to define the numbered rows and column groups.

**Fig. 4.**
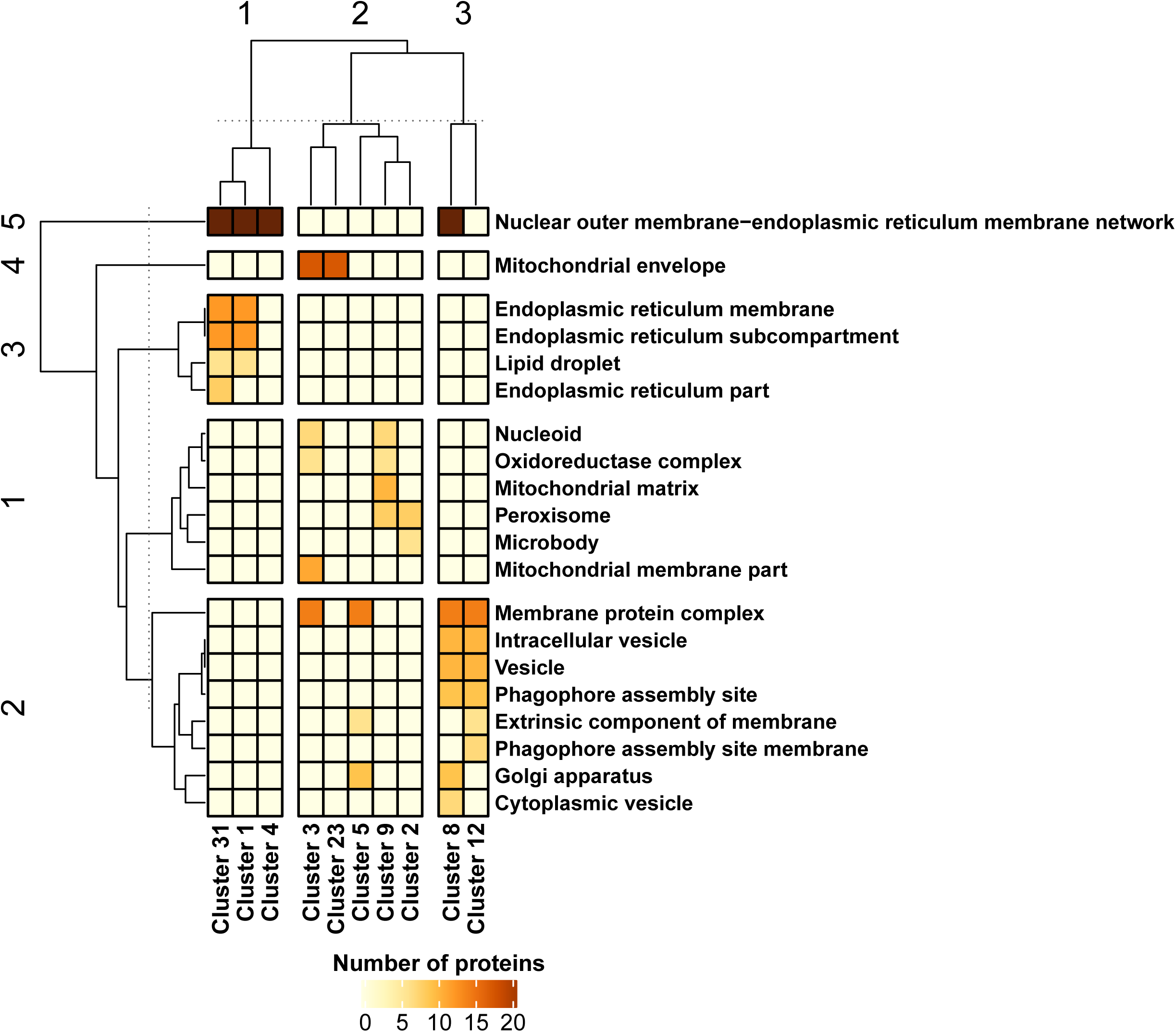
Heatmap showing the clustered cellular components based on the Gene Ontology (GO) analysis of autophagy, lipid metabolism, and proteostasis (ALP) network-associated communities/clusters. Heatmap rows and columns were grouped using the *k-*means distance-based method. Horizontal and vertical dotted lines indicate the cut-off point used to define the numbered rows and column groups.

### 3.4 Evaluation of ALP Pan-DEGs linked to LD structure and function

Using the available data on the various proteins associated with LD (Grillitsch et al., 2011) and the ALP Pan-DEGs identified in this study, 17 ALP Pan-DEGs linked to LD structure were identified as being upregulated in lager yeast cells during beer fermentation compared to during yeast biomass propagation (Suppl. Fig. 5A and B; Table S11). Of these 17 LD-associated ALP Pan-DEGs, 11 are directly involved with lipid metabolism and six are related to proteostasis (Table S11).

This result was supported by a proteome and transcriptome multilayered chemical-protein interaction network of the 17 LD-associated ALP Pan-DEGs (LDP network; Fig. 5), where it could be observed that *ERG6, ERG27, POX1, FAA4, YJU3, GPT2, ERG1, DPL1*, and *PDI1* were found to be overexpressed in transcriptome and also in publicly available yeast proteolipidome data from Casanovas et al. (2015) (Fig. 5 and Table S12). Additionally, the LDP network also contains lipid metabolism-associated nodes that were exclusively found to be expressed in proteolipidome data, like *TGL4, FAA1, FAS1, ERG4, ERG7*, and *ERG9*, which are connected with LD-associated ALP Pan-DEGs (Fig. 5 and Table S12).

**Fig. 5.**
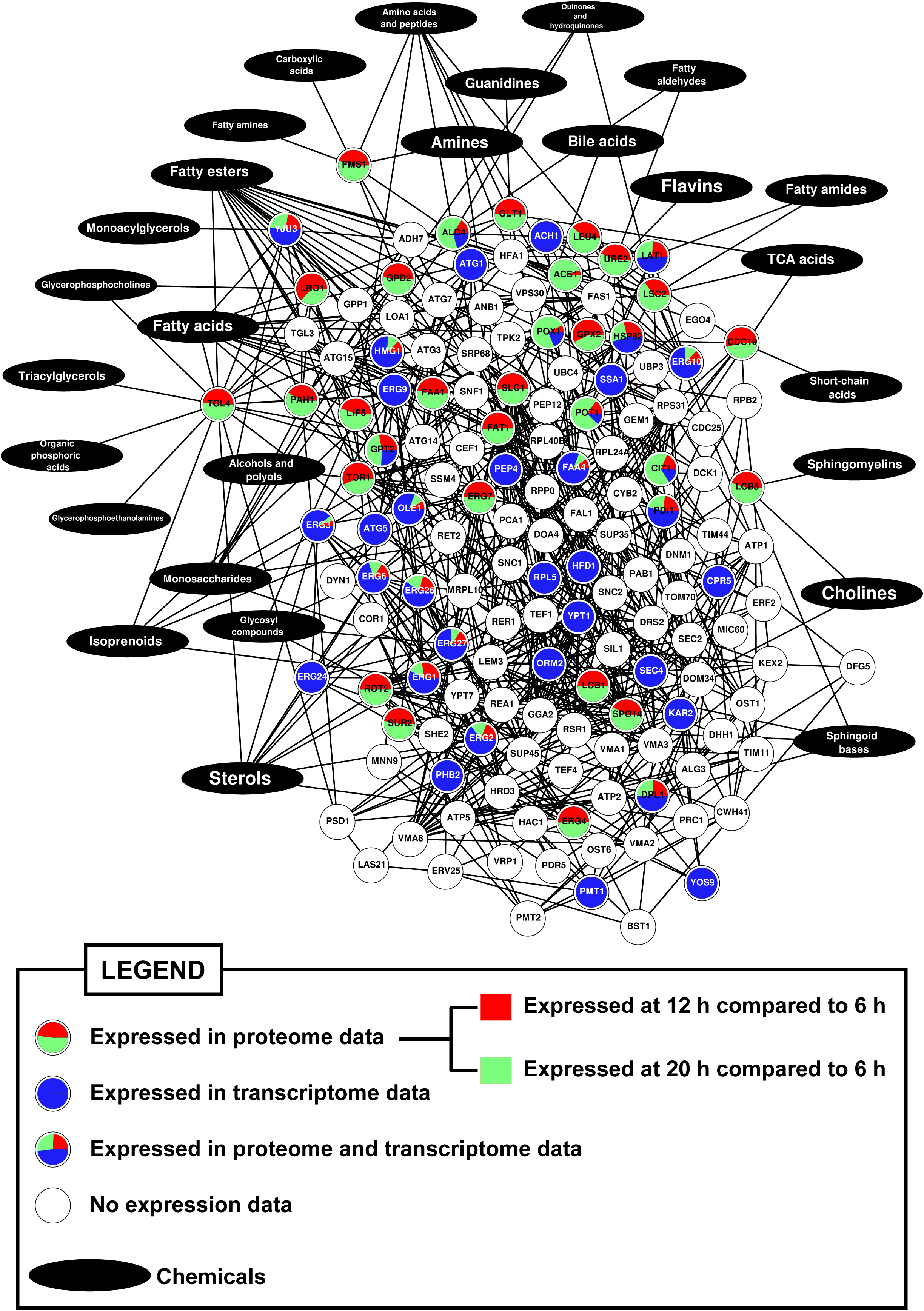
Shortest-pathway chemical-protein interaction (CPI) network of lipid droplets-associated proteins and molecules linked to lipid metabolism (LDP network). Expression data from proteome (Casanovas et al., 2015) and transcriptome (this study) analyses were added to network and are indicated as pie charts inside the nodes in different colors. Each pie slice is proportional to proteome and/or transcriptome expression values for a given node (please see Table S12 for additional information).

## 4. Discussion

The transcriptome and systems biology results obtained in this study suggest that, during lager beer fermentation, various ALP-associated genes are upregulated and their products cooperate in a shortest-pathway PPI network that is composed of multiple communities of proteins (Fig. 2A and D) in order to deal with the nutritional and ethanol stress observed during beer fermentation (Gibson et al., 2008, 2007). The GO analysis of these communities showed that they are associated with a large number of processes related to autophagy, mitochondria organization and activity, ER stress and Golgi organization, lipid/ergosterol metabolism, cytoplasmic vesicles, LDs, and phagophore assembly sites (Suppl. Fig. 3 and 4). All these processes are known to be modulated by the stressful conditions that yeast cells are exposed to during beer fermentation, including ethanol toxicity (Telini et al., 2020) and nitrogen and carbohydrate starvation (Gibson et al., 2007). Nitrogen starvation is a key condition that promotes autophagy in yeast cells (Cebollero and Reggiori, 2009), and despite this mechanism being well characterized in yeast strains during wine fermentation (Piggott et al., 2011), autophagy studies in lager yeasts are virtually absent. Interestingly, malt-derived wort used for beer fermentation contains different types of nitrogen sources that can prevent the activation of autophagy (Gibson et al., 2007); however, many autophagy-associated genes are upregulated during wine fermentation even in the presence of nitrogen sources (Piggott et al., 2011). In this study, the transcriptome data indicated the upregulation of various autophagy-related (*ATG*) genes during lager beer fermentation (Table S10). Many of these genes are involved in the regulation of macroautophagy and phagosome formation, but they are also related to microautophagy of organelles such as mitochondria and nuclei (Table S10). Among the *ATG* genes that were upregulated during beer fermentation, three (*ATG1, ATG8*, and *ATG18*) were characterized as major HB nodes in the ALP network (Table S10). *ATG1, ATG8*, and *ATG18* are part of the so-called “autophagy core machinery”, which is important for both micro- and macroautophagy (Lynch-Day and Klionsky, 2010). The transcriptome and systems biology data indicate that, during beer fermentation, the cytoplasm-to-vacuole targeting (Cvt) mechanism, mitophagy, and PMN are potentially activated (Fig. 3), all three of which depend on the activity of Atg1p, Atg8p, and Atg18p (Kanki et al., 2015; Krick et al., 2008; Lynch-Day and Klionsky, 2010). Although the importance of Cvt, mitophagy, and PMN to brewing yeasts strains is unknown in the context of beer fermentation, it has been reported that all three of these microautophagy processes are important for yeast cell adaptation in fermentative environments (Cebollero and Gonzalez, 2006; Kurihara et al., 2012).

Another fundamental aspect of micro- and macroautophagy mechanisms that should be considered is the formation of autophagosomes, which are membranous vesicular structures that deliver various cargo components for degradation in vacuoles (Lamb et al., 2013). The formation of autophagosomes strongly depends on the biosynthesis of lipids such as triacylglycerols and sterol esters, which originate from ER membranes in the form of LDs (Velázquez et al., 2016). In fact, low levels of nitrogen sources and the presence of glucose in yeast stimulate lipogenesis and increase the number of LDs, which, in turn, are required for efficient autophagy (Li et al., 2015).

The transcriptome and systems biology data demonstrated the upregulation of several genes linked to neutral lipid and ergosterol biosynthesis in lager yeast cells during beer fermentation (Table S10). Moreover, the GO analysis of cellular components linked to clusters 1, 4, and 31 indicated associations with ER membrane structure, the ER association with the nuclear outer membrane, and LDs (Suppl. Fig. 3 and 4).

Besides the importance of lipid biosynthesis for autophagy, lipids may be key regulators of proteostasis and modulators of the permeability to ethanol of the cell membrane (Aguilera et al., 2006; Chi and Arneborg, 1999; Ma and Liu, 2010). In fact, lipid metabolism and LDs are important factors in proteostasis, as it has been shown that yeast cells exhibiting defective biosynthesis of neutral lipids and LDs have chronic ER stress (Graef, 2018; Velázquez et al., 2016). In addition to being important during proteostasis, LDs are essential for macroautophagy (Velázquez et al., 2016; Velázquez and Graef, 2016), helping to assemble autophagosomes and induce mitophagy (Carmona-Gutierrez et al., 2012) and microlipophagy (Vevea et al., 2015). Notably, six upregulated proteostasis-associated genes identified in this study that are linked to the inter-organellar proteostasis mechanism (*CPR5, KAR2, PDI1, PMT1, RPL5*, and *SSA1*) (Telini et al., 2020) are also found in LD structures (Suppl. Fig. 5B and Table S10). Interestingly, ER stress can induce the formation of LDs (Fei et al., 2009) and LDs interact with mitochondria and peroxisomes via Erg6p (Pu et al., 2011), a protein encoded by *ERG6*, which was found upregulated in this study (Suppl. Fig. 5B and Table S10). A supporting proteome and transcriptome multilayered chemical-protein interaction network (LDP network) using the proteolipidome data of Casanovas et al. (2015) pointed that many of the overexpressed LD-associated genes found from meta-transcriptome analysis are proteomically expressed during the transition of fermentation to respiration in yeast cells, including genes linked to synthesis of LD-associated fatty acids and triacylglycerols, like *FAA4* (Fig. 5 and Table S12). The fermentation-to-respiration transition is also observed in yeast cells during brewing, with the exception that the anaerobiosis and nutrient depletion in beer avoid the use of ethanol as carbon source by yeast cells (Gibson et al., 2007). Interestingly, Erg6p, Erg27p, Pox1, Faa4p, Yju3, Gpt2p, Erg1p, Dpl1p, and Pdi1p were also found to be differentially expressed in the proteome data of industrial wine yeast strains during wine fermentation (Rossouw et al., 2010). Additionally, Erg6p, Erg27p, Faa4p, and Yju3p were found in high resolution proteome analysis of purified LD fraction from yeast (Currie et al., 2014), also including Fat1p, Tgl3p, and Tgl4p, which are present in the LDP network (Fig. 5). Unfortunately, proteome data of brewing yeasts in condition of beer fermentation are still very limited; however, a proteome study of an ale yeast strain indicated that the many of overexpressed proteostasis- and lipid metabolism-associated genes found in this work (Suppl. Fig. 4 and Table S7; Fig. 5 and Table S12) were also found to be expressed in the proteome of ale yeast A38 strain during beer fermentation (Kobi et al., 2004). Thus, the systems biology data gathered in this work support the idea that beer fermentation promotes lipid biosynthesis in yeast cells, specially unsaturated fatty acids (UFAs) and ergosterol, which can be related to the composition of LD’s membrane. In fact, it has been reported that the levels of UFAs and ergosterol increased during fermentation in order to antagonize the membrane fluidity induced by ethanol (Ding et al., 2009) and LD-associated triacylglycerols and sterol esters can differentially change during the fermentation-to-respiration transition (Casanovas et al., 2015).

Finally, LDs have a central role in inter-organellar communication and the promotion of autophagy and proteostasis, and it is essential to determine how LDs are produced and regulated during beer fermentation. Moreover, it is essential to determine how LDs are produced and regulated in the context of a hybrid species such as the lager yeast *Saccharomyces pastorianus* (Gorter de Vries et al., 2019), in comparison to its parental counterparts (*Saccharomyces eubayanus* and *Saccharomyces cerevisiae*), in order to design new lager beer strains that are resistant to high ethanol concentrations during HG/VHG beer production.

## Compliance with ethical standards

### Funding

This work was supported by the Conselho Nacional de Desenvolvimento Científico e Tecnológico [grant number 302969/2016–0]. The sponsor had no role in the study design; collection, analysis, and interpretation of data; writing of the report; or decision to submit the article for publication.

### Declarations of interest

none

### Human and animal rights

The experiments included in this manuscript did not involve any animal or human participants.

## Notes

https://github.com/bonattod/Lipid_stress_data_analysis.git

### Competing Interest Statement

The authors have declared no competing interest.

